# High-dimensional phenotyping to define the genetic basis of cellular morphology

**DOI:** 10.1101/2023.01.09.522731

**Authors:** Matthew Tegtmeyer, Jatin Arora, Samira Asgari, Beth A. Cimini, Emily Peirent, Dhara Liyanage, Gregory Way, Erin Weisbart, Aparna Nathan, Tiffany Amariuta, Kevin Eggan, Marzieh Haghighi, Steven A. McCarroll, Anne E. Carpenter, Shantanu Singh, Ralda Nehme, Soumya Raychaudhuri

## Abstract

The morphology of cells is dynamic and mediated by genetic and environmental factors. Characterizing how genetic variation impacts cell morphology can provide an important link between disease association and cellular function. Here, we combined genomic and high-content imaging approaches on iPSCs from 297 unique donors to investigate the relationship between genetic variants and cellular morphology to map what we term cell morphological quantitative trait loci (cmQTLs). We identified novel associations between rare protein altering variants in *WASF2, TSPAN15*, and *PRLR* with several morphological traits related to cell shape, nucleic granularity, and mitochondrial distribution. Knockdown of these genes by CRISPRi confirmed their role in cell morphology. Analysis of common variants yielded one significant association and nominated over 300 variants with suggestive evidence (P<10^-6^) of association with one or more morphology traits. Our results showed that, similar to other molecular phenotypes, morphological profiling can yield insight about the function of genes and variants.

## Introduction

Cellular morphology is an important and informative cellular trait across cell types, health, and disease. Changes in cell morphology can be indicators of disease. A classic example is sickle cell anemia, which gets its name by the sickle-like morphology of blood cells observed in patients afflicted with this condition^1^. Like other traits, such as gene expression, cellular morphology is, in part, genetically determined. Genetic studies have implicated various loci associated with red blood cell phenotypes such as mean volume and hemoglobin content^2,3^. However, there is still limited understanding of how human genetic diversity shapes cell morphology. This is due to several challenges: cell morphology is hard to quantify; ascertaining how human genetic variation influences cellular phenotypes in living biological systems can be challenging and cost prohibitive; and many cellular phenotypes are context-specific requiring the acquisition of relevant tissue and cell types. Profiling cell morphology in different cell types and across genetically diverse populations could facilitate the identification of such loci.

Recent innovations in cellular imaging and analysis have made it possible to measure thousands of morphological traits from a single cell, constructing morphology based ‘profiles’. Cell Painting, for example, leverages multiplexed dyes to enable trait measurement across many cellular compartments and organelles^4,5^. Cell Painting can ascertain gene function by linking expression to cellular traits^6^ and has been used to enable the prediction of functional impacts from lung cancer variants^7^. Cell morphology profiling provides a great asset for functional genomics studies compared to methods such as gene expression as it’s much more affordable and easily scalable at the bulk and single cell level. We hypothesized this approach could be leveraged to elucidate relationships more broadly between cell morphology and genetic variants.

Ascertaining how human genetic diversity influences cellular phenotypes in living biological systems has been difficult. Collections of induced pluripotent stem cells (iPSCs) provide a powerful tool for modeling human genetics^8^. The emergence of large iPSC collections, now available in several public repositories, provides access to cell lines from donors of diverse ancestry and genetic backgrounds, enabling the study of how human common and rare genetic variation impacts cellular function and behavior^9–17^. Attempts to investigate how genetic variants drive cell morphology using iPSC-based models have shown promise but have been restrained by insufficient sample size and by the limited number of cellular traits being measured, hampering discovery potential^18^.

Here, we identified the morphological impacts of genomic variants, or cmQTLs, by generating high-throughput morphological profiling and whole genome sequencing data from 297 unique cell lines. Using Cell Painting on >5 million iPSCs derived from these donors, we comprehensively quantified 3,418 cell morphological traits and assessed associations with rare and common genetic variants genome-wide. We identified trait-associations with rare-variant burden in several genes including *WASF2, PRLR*, and *TSPAN15* which we then functionally validated using CRISPR interference. Additionally, we found only one common variant convincingly associated with morphology but found suggestive evidence for over 300 loci. These findings show that similar to gene expression, the morphology of cells is mediated by genetic determinants and highlights the utility of image-based methods for functional genomics.

## Results

### Whole-genome sequencing and morphological profiling for 297 unique iPSC lines

To study associations between genetic variants and morphological traits, we assembled a cohort of iPSC lines from 297 unique donors, for which we had sex, ancestry, and clinical diagnosis information (**Fig. 1, Table S1, Fig. 2A**). We performed 30X whole-genome sequencing (WGS) on all iPSC lines. Following quality control (QC, see **Methods**), we retained 7,020,633 common (minor allele frequency (MAF) > 5%) and 122,256 rare (MAF < 1%) variants for downstream analyses.

**Fig 1.**
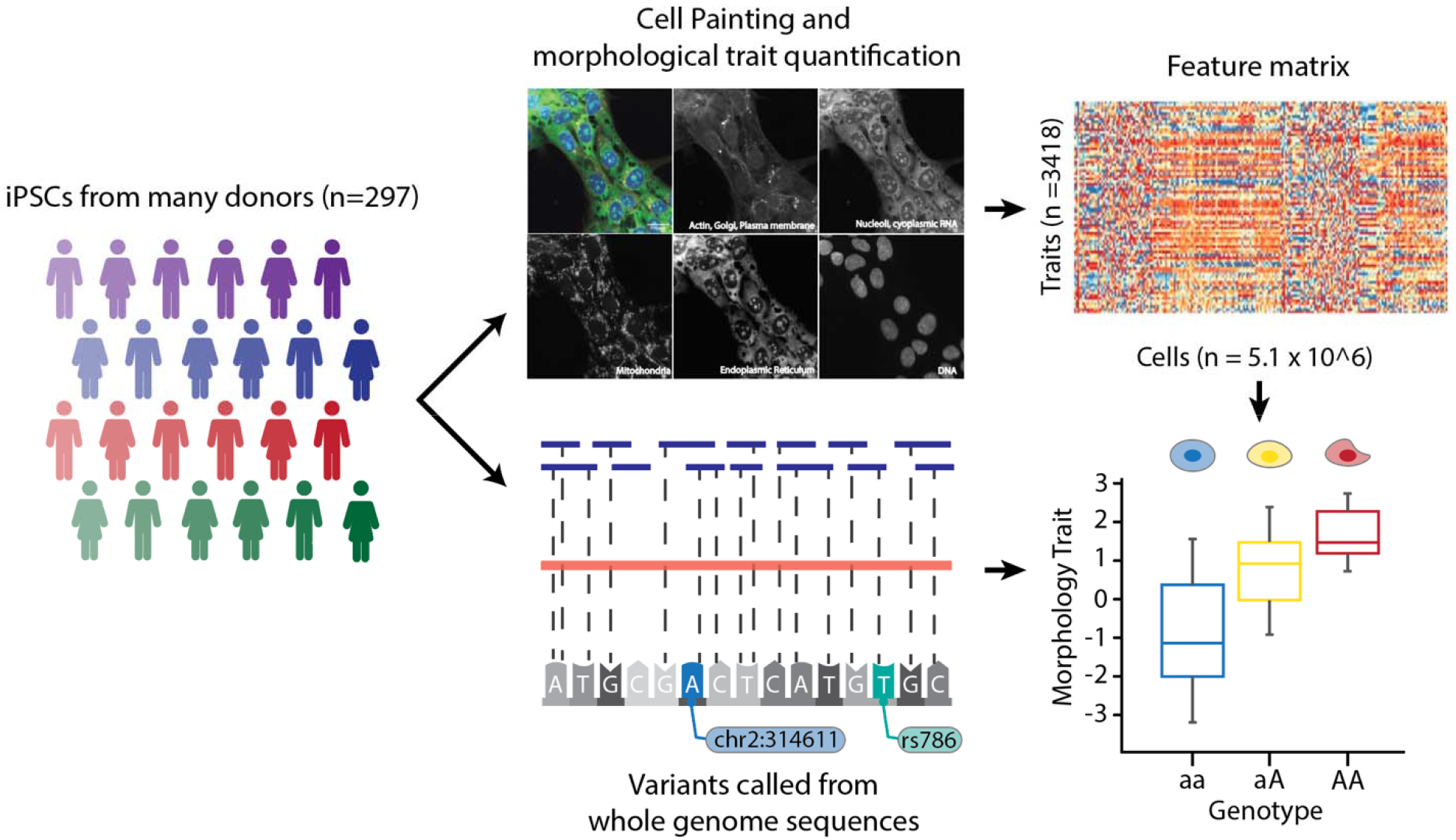
Study Overview. iPSC lines from 297 donors were expanded, quality-control checked and then subject to both high-throughput imaging with Cell Painting and 30X whole-genome sequencing. Overall, we imaged 5.1×10^6^ cells across all donors and quantified 3,418 morphological traits per cell using CellProfiler software. We inferred genetic variants from the WGS data and investigated whether individual morphological traits associated with both rare and common variation.

**Fig 2.**
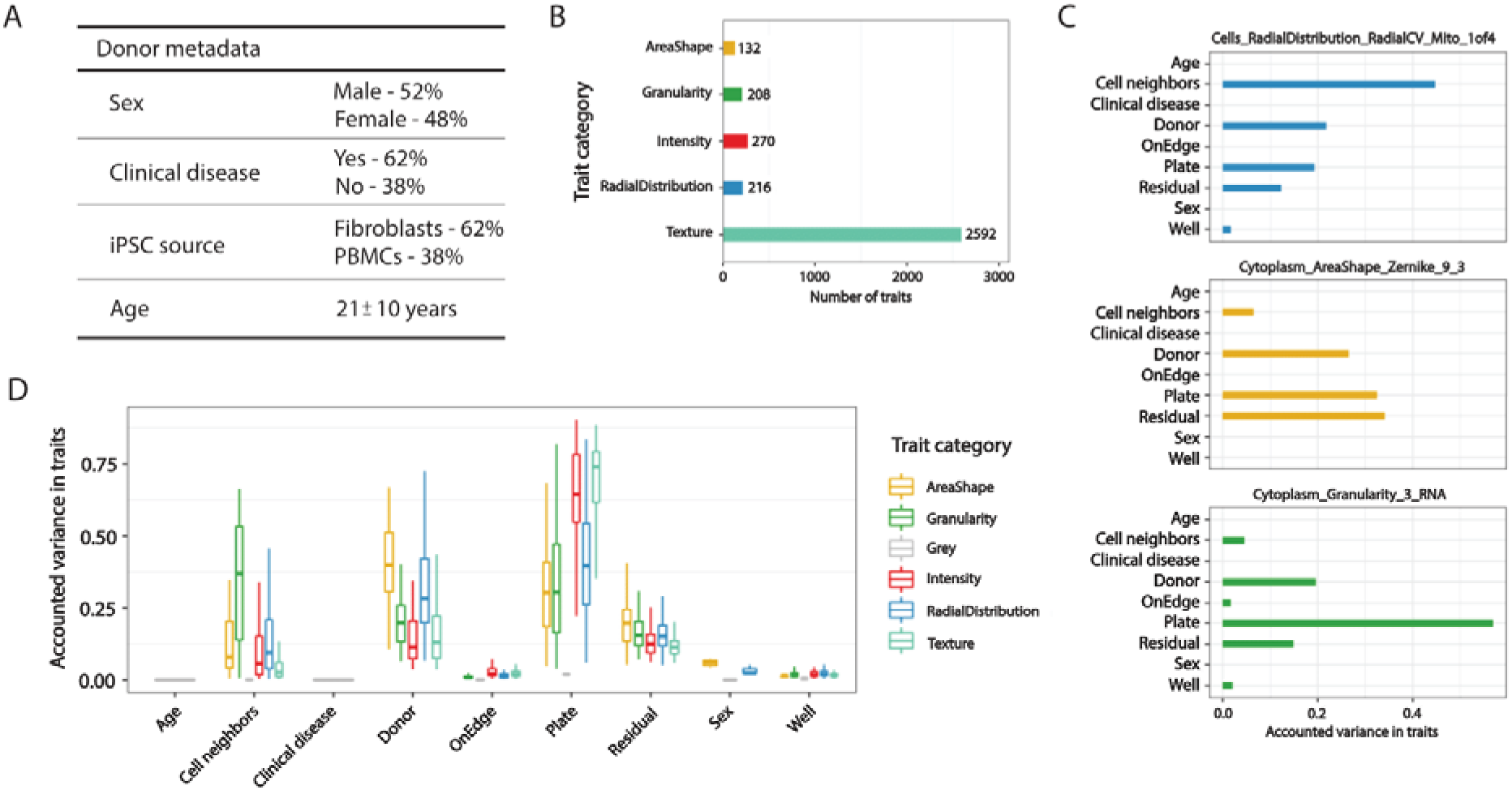
Summary of morphological traits and variant component analysis. (**A**) Table with cell line metadata (**B**) Summary of five categories of morphological traits captured in our data (n=3418) (**C**) Exploring explained Variation in individual traits, namely distribution of mitochondria around nucleus, cytoplasmic Zernike shape metric 9_3, and cytoplasmic granularity in the RNA channel at scale 3, showed differences in sources of variance, including technical effects such as plate and well position and biological sources such as donor. Donor ID represents the remaining difference among profiles after accounting for all other technical, demographic, and disease-status metadata. (**D**) Same as C but for all morphological traits (n=3418)

To quantify cellular traits, we adopted the Cell Painting assay for use across a large array of different iPSC lines. This multiplexing dye assay uses six stains to capture morphological characteristics for eight cellular compartments including mitochondria, cytoplasmic RNA, actin cytoskeleton, nucleoli, endoplasmic reticulum, Golgi apparatus, plasma membrane, and nuclei^4,5^.

Overall, we measured 3418 morphology traits for 5.1 million iPSCs from 297 donors after stringent QC (**Methods, Fig. S1**, **Table S2**). We classified all morphological traits based on the cellular characteristics they represented, yielding five categories: Area and shape, Granularity, Intensity, Radial distribution, and Texture (**Fig. 2B**).

### Principal components and variance component analyses

To assess if cell morphological may have a genetic component (cell morphological quantitative trait loci; cmQTLs), we assessed if replicates are correlated after correcting technical factors such as plate and well batch effects. These factors have previously been shown to alter morphology-based readouts^19^. Additionally, we explored how demographic factors including donor sex, disease status, age at sample generation, and iPSC sample source tissue may contribute to these traits (**Fig. 2A**). We observed non-random segregation of iPSC lines in principal component analysis (PCA) of morphology traits across ancestry categories (**Fig. S2**) and across plates (**Fig. S3**), indicating the contribution of genetic and technical factors to the measurement of morphology traits. To identify and control for these factors, we generated per-well pseudo-bulk trait profiles through mean-averaging of single cell profiles, resulting in eight measurements per trait per donor, one for each of the eight replicates. With our pseudo-bulked well-level data, we performed variance component analysis to quantify the observed variance that can be attributed to each morphological trait (**Methods**). We assessed the significance for each variance component, correcting for the number of tests, which was the product of the traits (*n*=3418) and factors (*n*=9, namely iPSC cell line, plate and well of sequencing, whether the well was on the plate edge, tissue of origin for iPSC cell line, average number of cell neighbors (other cells in contact with a given cell) in the well, donor’s sex, age, and disease status). Plate effects were associated with 3417 traits and explained 61.8+17% of the variance, thus having a major impact on morphology (**Fig. 2D**). We found several confounders which contributed varying levels of effect on different morphological traits (**Fig. 2C**). After correcting for these covariates, 16.7+11% of variance in all morphological profiles was explained by cell line donor, indicating the potential for a genetic basis to the variability in morphology traits (**Fig 2D**). Interestingly, the difference among donors explained a greater degree of variance in the trait category of AreaShape relative to the other trait categories (Wilcoxon rank sum test *P* = 1.1×10^-55^, **Fig. S4**). We note that some of the shared variance may be explained by non-genetic factors, such as stable epigenetic modifications.

### Selection of traits for association analysis

We next summarized well-level trait values into donor-level values (i.e., pseudo-bulk) by mean-averaging individual traits across all wells per donor, resulting in one measurement per trait per donor (N = 3418 traits and 297 donors). As cells often display varied morphology in response to environmental cues, we segregated all cells into two groups based on whether they had any cells in contact (called colony cells, 97.48% of all cells) or not (called isolate cells, 2.52% of all cells)^18,20^. In both colony and isolated cells, most traits (93.7 and 91.2%, respectively) had very high pairwise correlation (*Pearson r* > 0.9) with one or more traits (**Fig. S5**), suggesting the presence of many traits that were not independent of each other. Therefore, to reduce redundancy in association analysis, we examined pairwise correlation (*Pearson r*) among all 3418 morphological traits across colony and isolated cells and selected a common set of 246 traits having *r* < 0.9 with each other by iteratively selecting a single representative trait for the set of correlated traits (*r* > 0.9) (**Methods**). We refer to this common set of 246 traits as “composite traits” **(Table S3).**

### Rare variant association analysis

We next explored the effect of rare genetic variation on cellular morphology. We investigated the association of composite traits (*n* = 246) with gene-level burden of protein-altering rare variants (MAF < 0.01). To ensure well-powered investigation, we only examined 9105 genes that had rare variants in at least 2% of donors (*n* >= 6) for our association analysis. We modeled individual morphology traits as a function of rare protein-altering variant burden in a gene, controlling for plate, well, and donor sex using linear regression (**Methods, Fig. S6**). We performed this analysis separately in colony and isolated cells. Of all tested traits, one trait in colony cells and 3 traits in isolated cells passed the genome-wide significance threshold (*P* < 2.2×10^-8^, Bonferroni correction for 246 traits and 9105 genes) (**Fig. 3A**). We did not observe any inflation in association statistics for these traits (Lambda (λ) = 1.01 for the association in colony cells and λ = 1.01, 0.96, 0.98 for the associations in isolated cells) (**Fig. S7**).

**Fig 3.**
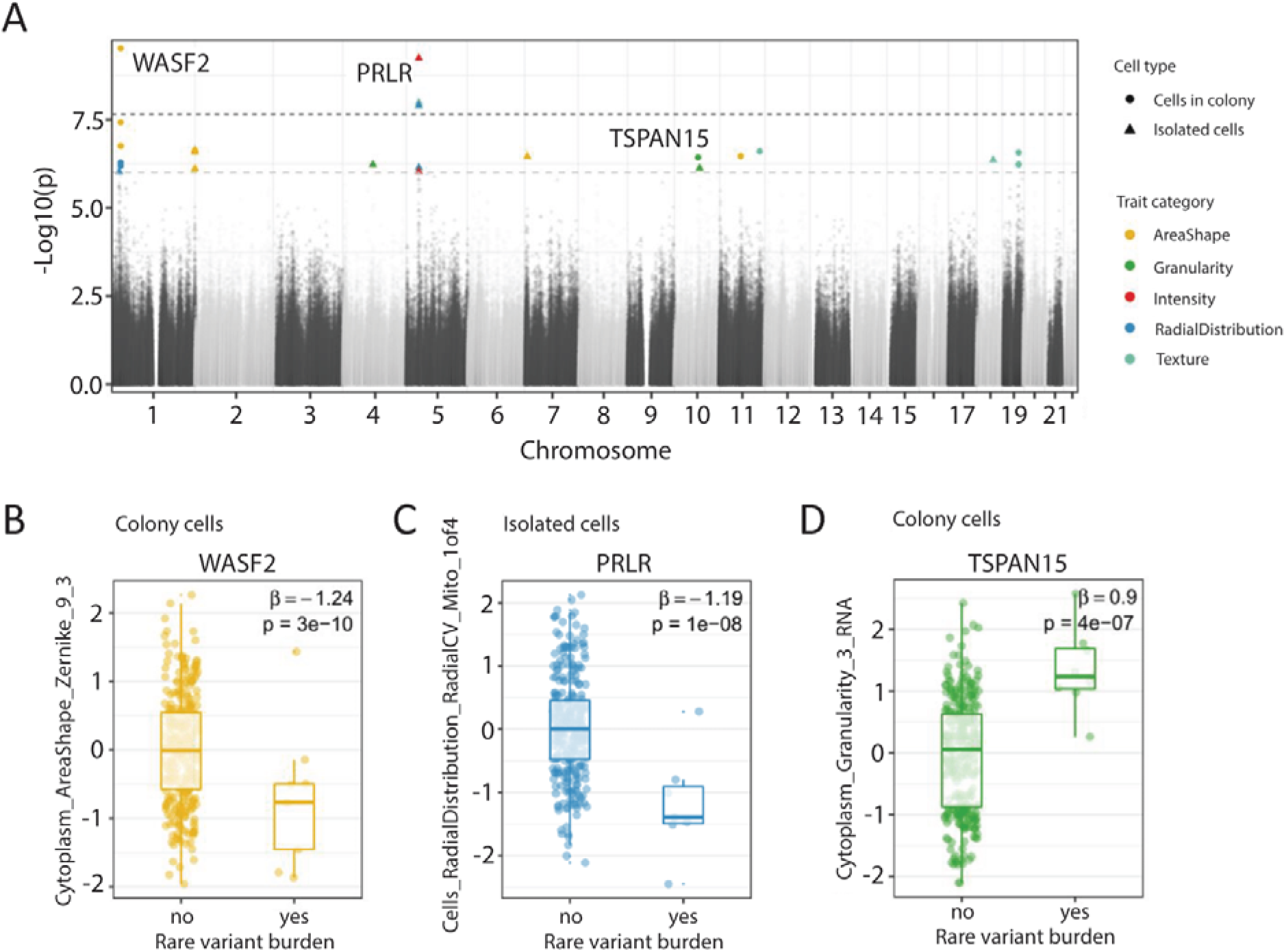
Association between morphology and rare variant burden. (**A**) Manhattan plot showing association between morphological traits (n=246) and rare variant burden in candidate genes (n=9105). Black dotted line represents the p-value threshold after Bonferroni correction for the number of tested traits and genes (*P* = 0.05/246×9105, i.e. 2.2×10^-8). Grey dotted line represents the p-value threshold for suggestive evidence of association (*P* = 10^-6). (**B-D**) Box plots displaying the association between the Zernike_9_3 cytoplasm shape metric and rare variant burden in *WASF2* gene (B), distribution of mitochondria around the nucleus and rare variant burden in *PRLR* gene (C) and cytoplasmic granularity measure in the RNA channel and rare variant burden in *TSPAN15* gene. We provide the effect size (β estimate) and raw p-value of the association for each trait.

In colony cells, a Zernike shape measure of the cytoplasm (*Cytoplasm_AreaShape_Zernike_9_3*) was negatively associated with rare variant burden in the *WASF2* gene (*n* = 3 missense and 1 in-frame deletion rare variants, β or effect size (se) = −1.24 (0.18), *P* = 3.1×10^-10^; **Fig. 3B**). *WASF2* protein binds profilin, a G-actin-binding protein, promoting the exchange of ADP/ATP on actin and the formation of actin filament clusters^21,22^. The disruption of *WASF2* impairs actin formation and organization that could lead to their polarized distribution and spindle-shaped cells^23^. In representative images of cells with rare variants in *WASF2* it is difficult to identify this polarized and spindle-like shape by eye when compared to reference lines (**Fig. S8**). As many phenotypes may only be uncovered using analyses such as these, it highlights the necessity of leveraging high-dimensional morphological profiling over more traditional methods of capturing cellular phenotypes. Moreover, rare variant burden in *WASF2* had nominal association (*P* < 0.05) with 90 other traits including 27 traits of area and shape category, substantiating *WASF2* as a genetic determinant of cellular morphology **(Table S4)**.

In isolated cells, three traits were associated with rare variant burden in the *PRLR* gene, one of which was the asymmetry in the distribution of mitochondria in the perinuclear space (*Cells_RadialDistribution_RadialCV_Mito_1of4*, *n* = 6 missense rare variants, β (se) = −1.17 (0.2), *P* = 1.2×10^-8^; **Fig. 3C**). *PRLR* encodes membrane-anchored receptors for a prolactin ligand and is a part of the class-I cytokine receptor superfamily and regulator of JAK-STAT5 pathway activity^24^. In addition to its well-known role in pregnancy and lactation, *PRLR* also plays a key role in an autocrine/paracrine loop present in stem cells, mediating their quiescence and proliferation^25^. Previous findings in adipocytes showed *PRLR* KO alters mitochondrial packing and distribution throughout the cell^26^. In a mouse model of depression, silencing of the *PRLR* gene inhibited neuron apoptosis, suggesting that disruption of *PRLR* activity could lead to cellular proliferation^27^. Indeed, we observed a higher cell count in iPSC lines carrying a rare variant burden in the *PRLR* gene compared to reference iPSC lines (**Fig. S9**). Further evidence supports a link between mitochondria distribution and neurodegeneration within the aging brain, whereby the position of mitochondria with respect to different organelles is essential for supplying bioenergetic homeostasis to cellular compartments^28–32^. These findings suggest mutations in *PRLR* drive asymmetry of mitochondria within the perinuclear ring, improving the bioenergetic homeostasis of the cell’s nucleus and reducing cellular apoptosis. Moreover, rare variant burden in *PRLR* had nominal association (*P* < 0.05) with 118 other traits, providing more support to *PRLR* as a genetic determinant of cellular morphology **(Table S5)**.

We also inspected the associations with suggestive evidence, i.e., *P* < 10^-6^. There was a total of 12 and 13 associations in colony and isolated cells, respectively, which passed this threshold **(Table S6)**. Of our suggestive hits, one of the strongest associations was between the distribution in size of RNA particles in the cytoplasm (*Cytoplasm_Granularity_3_RNA*) and rare variant burden in *TSPAN15* gene (*n*=2 missense and 1 splice region rare variants in the gene, β (se) = 0.9 (0.17), *P* = 3.7×10^-7^; **Fig. 3D**). *TSPAN15* is expressed in all human tissues and encodes for a cell surface protein^33^. This protein plays a role in cell activation, development, and proliferation by negatively regulating Notch-signaling activity^34^, indicating that disruption of *TSPAN15* could lead to higher transcriptional activity and RNA amount in the cell proxied by higher cytoplasmic RNA granularity and cellular proliferation. Indeed, all iPSC lines carrying a rare variant burden in *TSPAN15* had higher cell count compared to wild-type iPSC lines (**Fig. S10**).

To ensure that the observed associations were not driven by somatic variation potentially introduced during iPSC generation, cell seeding or genome sequencing, we repeated our analysis restricting to only those variants that were previously observed in the gnomAD database^35^ (106,590 of 122,256 variants). We recapitulate all observed associations (significant after Bonferroni correction for multiple testing and with suggestive evidence) with concordant effect size and statistical significance (p-value) (**Fig. S11**). Taken together, our analyses indicated that, using our dataset, we could successfully identify associations between rare coding variants and several morphological traits.

### Functional validation of rare variant associations

CRISPR-based gene editing has been shown to be a viable mechanism for validating gene expression phenotypes resulting from rare-variation^36^. To validate our rare-variant burden associations, we tested whether knockdown of these genes impacted the same traits for which we identified a rare-variant burden association. We transduced iPSCs from a cell line expressing constitutive dCas9-KRAB CRISPRi machinery with sgRNAs targeting *WASF2, PRLR*, and *TSPAN15*(**Fig. 4A**). We targeted each gene with 2 different sgRNAs, and validated each sgRNA for knockdown in expression of their gene target, showing a range of knockdown efficiency (15%-95%) (**Fig. S12, Table S7**). Cells transduced with sgRNAs were Cell Painted and morphological traits were extracted and quantified using the same pipeline from our discovery cohort.

**Fig 4.**
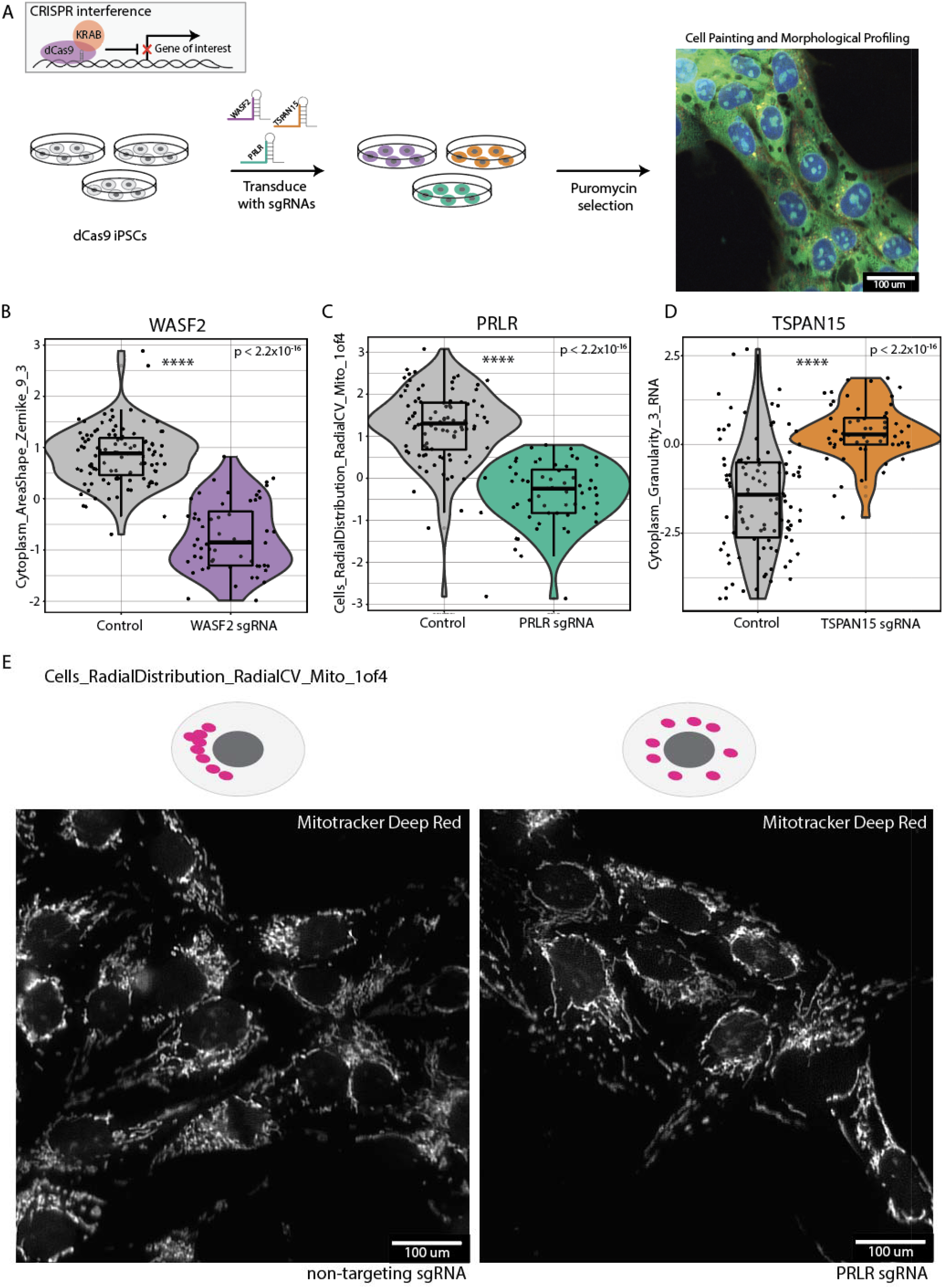
Functional validation of rare-variant burden associations. (**A**) Workflow for knockdown of rare-variant genes using CRISPR interference in iPSCs expressing constitutive dCas9-KRAB. (**B-D**) Violin plots displaying quantification of traits between control non-targeting sgRNAs and sgRNAs targeting *WASF2, TSPAN15*, and *PRLR* on a per-well level *(n* = 56 non-target sgRNAs, *n* = 56 targeting sgRNAs, *P* < 2.2×10^-16^, Welch’s Two-Sample T-Test) Effect on the trait score is consistent with what we observed in our rare-variant burden association. (**E**) Representative image of an observable gene-trait association for *PRLR*. Cells_RadialDistribution_RadialCV_Mito_1of4 relates to the asymmetric distribution of mitochondria in the ring right around the nucleus. In the non-targeting controls (left) we observed clustering of mitochondria on a particular side of the nucleus, whereas in the *PLRL* knockdown sgRNA (right) we observed a more distributed presence of mitochondria around the nucleus.

For each of the three genes tested, we detected the predicted changes in each individual trait, and the change was in the same direction as our association analysis relative to non-targeting sgRNA controls (*n* = 28 wells per targeting sgRNA and 52 wells per non-targeting sgRNA, *Welch’s Two Sample T-Test, P < 2.2×10^-16^*) (**Fig. 4B-D**). Specifically, knockdown of *WASF2* resulted in a decrease in normalized score for the trait *Cytoplasm_AreaShape_Zernicke_9_3* (**Fig. 4B**). We further observed that a reduction in the expression of *TSPAN15* coincided with an increase in trait score for *Cytoplasm_Granularity_3_RNA* (**Fig. 4D**). Finally, knockdown of *PRLR* expression decreased *Cells_RadialDistribution_RadialCV_Mito_1of4*, which defines the relationship between the radial distribution of mitochondria around the nucleus (**Fig. 4C**). This effect is highlighted in representative images, whereby cells transfected with a *PRLR* targeting sgRNA display more uniform distribution of mitochondria around the nucleus when compared to non-targeting sgRNA cells where mitochondria tend to colocalize to one side of the nucleus (**Fig. 4E**).

### Common variant association analysis

To identify common variants that mediate cell morphology, we performed 246 genome-wide association analyses, one for each composite trait. Each association was tested in colony and isolated cells separately (**Fig. 5A, B**). With our set of 297 donors, only one variant, rs315506, overlapping the chr17q11.2, passed the genome-wide significance threshold (Bonferroni correction for 246 morphology traits, 5×10^-8^/246 = 2×10^-10^). rs315506 is an intergenic variant and was associated with spatial distribution of endoplasmic reticulum (ER) in the cytoplasm (*Cytoplasm_RadialDistribution_RadialCV_ER_3of4*) in colonies (MAF = 0.08, β (se) = −0.52 (0.08), *P* = 1.4×10^-10^, **Fig. 5C**). This variant also showed suggestive evidence of association (*P* < 10^-5^/246 = 4.1×10^-8^) with spatial distribution of ER near the periphery of cells (*Cells_RadialDistribution_MeanFrac_ER_4of4*). rs315506 lies in the center of a 400kb window containing the genes *NF1, CORPS, UTP6* and *SUZ12*. Microdeletions on chr17q11.2 cause NF1 microdeletion syndrome, which has been shown to impair protein localization to the ER^37,38^.

**Fig 5.**
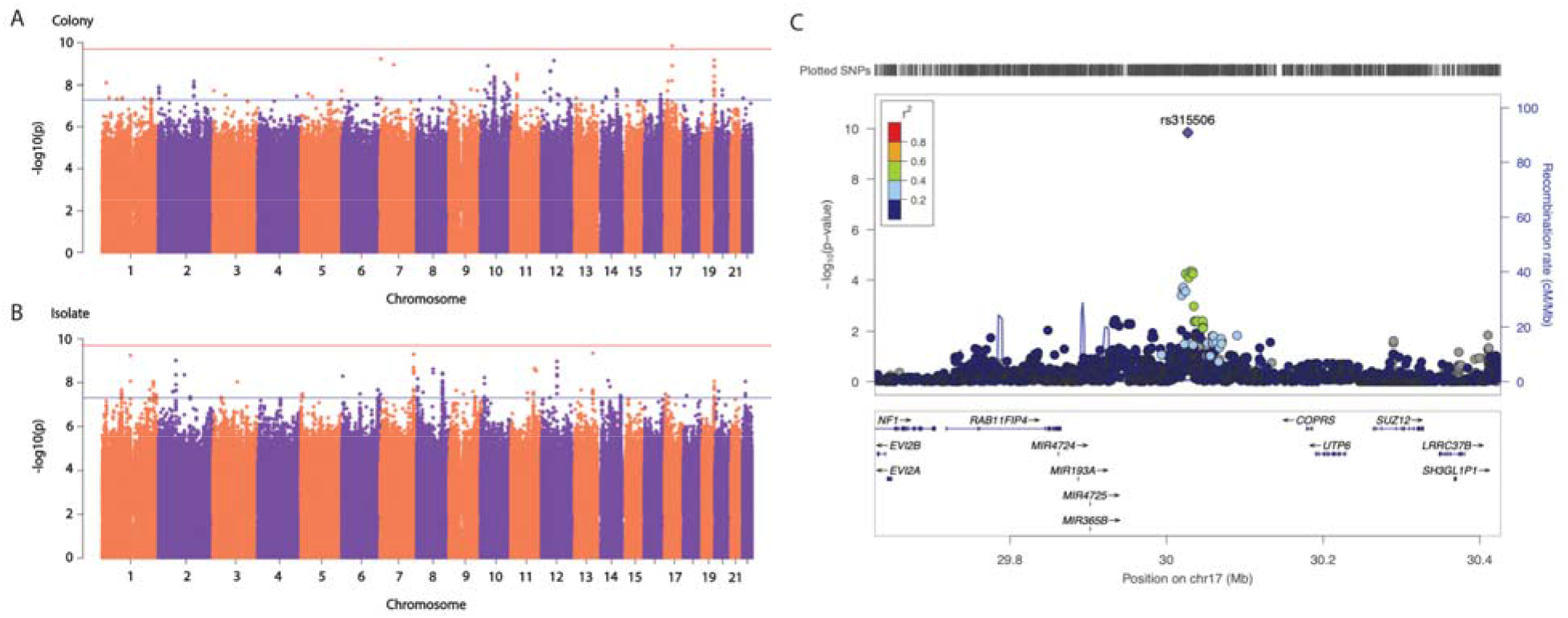
Common variant analysis. (**A**) Manhattan plot for trait association test in colony cells. Red line represents the p-value threshold after Bonferroni correction for the number of tested traits and genes (P < 2×10^-10^) Blue line represents the p-value threshold for hits with suggestive evidence (*P* < 4.1×10^-8^) (**B**) Manhattan plot for trait association test in isolate cells. Red line represents the p-value threshold after Bonferroni correction for the number of tested traits and genes (P < 2×10^-10^) Blue line represents the p-value threshold for hits with suggestive evidence (*P* < 4.1×10^-8^) (**C**) LocusZoom plot for the association signal at chr17q11.2. rs315506, was significantly associated with spatial distribution of cytoplasm (Cytoplasm_RadialDistribution_RadialCV_ER_3of4) in colonies (MAF = 0.08, effect size (se) = −0.52 (0.08), *P* = 1.4×10^-10^)

Besides rs315506, in colony cells, the second strongest association was on chromosome 7 (between *Nuclei_Granularity_9_AGP* and rs36036340, MAF = 0.08, β (SE) = 0.38 (0.06), *P* = 6×10^-10^). rs36036340 lies within the gene *PRKAR1B*. Variants in *PRKAR1B* have been linked to neurodevelopmental disorders and activity of *PRKAR1B* has been shown to regulate tumorigenesis^39–41^. *PRKAR1B* mediates PI3K/AKT/mTOR pathway signaling through direct interactions between *PRKAR1B* and PI3K-110alpha^39^. Given that MTORC1 activity is heavily influenced by the Golgi apparatus, and regulates cellular proliferation and cell cycle, variants in rs36036340 may impact PI3K/AKT/mTOR signaling, altering cellular morphology pertaining to the Golgi, actin, and plasma membrane.

In isolated cells, the most significant association was found on chromosome 13 (between *Nuclei_RadialDistribution_RadialCV_Brightfield_2of4* and rs9301897, MAF = 0.13, β (se) = −0.31 (0.05), *P* = 4.5×10^-10^). rs9301897 lies within the gene *GPC6*. Genetic variants in *GPC6* have been implicated in Alzheimer’s disease (AD), and TD43-proteinopathy, a hallmark of amyotrophic lateral sclerosis (ALS) pathology, has been shown to regulate *GPC6* activity^42,43^. *GPC6* and other glypicans are known to play a role in cell growth and cell division through cell surface receptor activation^44,45^. As nuclear movement within the cell is heavily regulated by cell cycle dynamics, variants in *GPC6* may alter nuclear localization by impacting mitosis.

In total, over 300 loci reached the suggestive genome-wide significance threshold (*P* < 4.1×10^-8^, **Table S8**) suggesting that a larger sample size and improved statistical power would be able to identify additional common variants associated with cell morphology. Moreover, several loci (**Table S8**) showed suggestive association with more than one trait suggesting shared genetic etiology among different morphological traits.

## Discussion

Previous studies linking genetic variants to cellular function have largely focused on human genes and alleles which mediate molecular phenotypes, such as gene or protein expression and chromatin accessibility^46–49^. Expanding on these studies, here we combine high-throughput cell culture techniques with cost-effective and high-dimensional image-based cell profiling (i.e., Cell Painting) to connect genetic variants to their morphological function in 297 donors.

Our work provides the largest to date exploration of genetic influences on cell morphology (what we term cmQTLs). Where previous studies have been limited by both sample size and the scale of morphological measurements, we combined whole genome sequence analysis with Cell Painting to define relationships between genetic variants and 3418 morphological traits extracted from >5M iPSCs. Leveraging these advances, we identified novel associations between rare variant burdens in the *WASF2, PRLR, TSPAN15* and cell morphological traits related to the cytoplasmic area and shape, nucleic granularity, and the distribution of mitochondria around the nucleus. These associations were validated by mechanistic information about these genes from the literature as well as CRISPR-mediated knockdown in our study.

In our common variant analysis, we found one significant result and 300 potential associations, indicating that future studies with larger sample sizes may help in elucidating other such loci. Interestingly, we observed no overlap in traits and associated variants between colony and isolated cells, suggesting a differential effect of genetic variation based on the environmental context of the cells. This is consistent with previous studies that have shown that intrinsic properties of cells may only come to light in the context of altering the cellular environment^18,20^. Further, we identified confounding factors that drive variation in cellular phenotypes which are important to address when performing similar studies. In particular, uniform cell densities across conditions is critical in imaging-based assays. To address this challenge, we incorporated automated liquid handling devices to reduce the latency of manually pipetting into 384 well microplates.

Our work has several limitations that highlight directions for future research. First, the cell types utilized in this study are in a basal, undifferentiated state. It will be valuable for future studies to explore these associations in more physiologically relevant contexts, where disease-associated variants are enriched^50^. These findings suggest this framework could be applied to relevant cells and tissues such as iPSC-derived differentiated cells, post-mortem brain samples or excisable somatic cells. Second, though our study provides the largest (to our knowledge) image-based iPSC phenotyping dataset, we are still underpowered to detect a significant number of high confidence common variant cmQTLs. Future efforts may require cross-institutional collaborations to adequately scale *in vitro* sample sizes for common variant cmQTL identification.

This approach holds significant promise for future studies leveraging human-derived, disease-relevant cell types for modeling the impact of genetic variation on cellular function. The use of imaging to capture phenotypes is particularly attractive in experimental designs for several reasons, such as the low cost per cell for imaging, and the ease of processing data and preparation of the cells or tissues as compared to the generation of other molecular data such as RNA-sequencing or epigenomic assays^51^. Moreso, large imaging datasets provide tools for developing robust statistical models for combined analysis of morphological profiling data with additional modalities such as gene expression to comprehensively interrogate genetic variants and their function^52^. Taken together, we demonstrate cellular morphology can be a cost-effective readout for modeling the biological function of human genetic variation.

## Supplementary Materials

### Online Methods

#### Material and iPSC generation

Our dataset comprised 297 donors from the iPSC repository of California Institute for Regenerative Medicine (CIRM) (Supplementary Table 1). Either B cells or Fibroblasts were taken from each donor from which iPSC lines were generated using non-integrating episomal vectors. Cells were cultured in StemFlex (ThermoFisher; cat#A334901) culture media and passaged for expansion with 1mM EDTA (Gibco; cat#15575020) and 10uM Y-27632 (StemcellTech; cat#72304). Each iPSC sample underwent Global Screening Array (GSA) for karyotype analysis to ensure chromosomal integrity, as well as 30X whole-genome sequencing to determine genome-wide variants for each donor. Each iPSC line was cultured between passages 12 and 15 before use in the experiment.

#### Cell seeding and staining

For each batch of imaging, cells were detached from 6-well NUNC plates using Accutase (StemcellTech; cat#07920) for generating single-cell suspensions. Following detachment, cells were centrifuged at 1000 rpm x 5:00 and re-suspended in StemFlex medium supplemented with ROCK inhibitor. After each cell line was counted to determine cell solution concentration and viability, the desired cell solution volume was aliquoted into a 96-deep well low attachment plate. To disperse a high number of cell lines across a 384-well plate in a semi-random fashion, we optimized the use of an Agilent Bravo liquid handling device. Here, using an 8-channel head, cell solutions were transferred from the 96-well low attachment plate and distributed into a geltrex-coated Perkin Elmer Cell Carrier 384-well plate.

#### Cell Painting and imaging

Cells were staining and imaged with minor adaptations to procedures described previously^4,5^. Six hours post seeding in 384-well plates, cells were treated for 30 min with 0.5 uM MitoTracker Deep Red FM - Special Packaging (Thermo Fisher cat#: M22426) dye at 37°C. Following the MitoTracker treatment, cells were fixed with 16% paraformaldehyde diluted to a final concentration of 4% (Thermo Fisher cat#: 043368.9M) for 20 minutes in the dark at RT. After three washes with 1X HBSS cells were permeabilized and stained using a solution of 1X HBSS (Thermo Fisher cat#: 14175095), 0.1% Triton-X-100 (Sigma Aldrich cat#: X100-5ML), 1% Bovine Serum Albumin, 8.25nM Alexa Fluor 568 Phalloidin (Thermo Fisher cat#: A12380), 0.005mg/ml Concanavalin A, Alexa Fluor 488 Conjugate (Thermo Fisher cat#: C11252), 1ug/ml Hoechst 33342, Trihydrochloride, Trihydrate (Thermo Fisher cat#: H3570), 6uM SYTO 14 Green Fluorescent Nucleic Acid Stain (Thermo Fisher cat#: S7576), and 1.5ug/ml Wheat Germ Agglutinin, Alexa Fluor 555 Conjugate (Thermo Fisher cat#: W32464) for 1hr at RT in the dark. Following the staining, plates were washed 3X with 1X HBSS and sealed until imaging. Cell Painted plates were imaged on a Perkin Elmer Phenix Automated Imaging system under a standardized protocol. All 297 cell lines were dispersed across seven plates which were imaged in four separate batches.

#### Quantification of cellular morphology traits and their quality control

The segmentation of individual cells in the image into its cellular compartments (whole cell, cytoplasm and nuclei) and subsequently quantification of morphology traits for each cellular compartments was done using CellProfiler 3.1.8^53^; pipelines are available at https://github.com/broadinstitute/imaging-platform-pipelines/tree/master/cellpainting_ipsc_20x_phenix_with_bf_bin1. Analysis of CRISPR experiments was done in CellProifler 4.2.4 with pipelines availalbe at https://github.com/broadinstitute/imaging-platform-pipelines/tree/master/cellpainting_ipsc_20x_phenix_with_bf_bin1_cp4^54^. Subsequently, cells missing measurement for more than 5% of traits were removed. Morphology traits *a priori* known to be problematic, not measured across all cells or non-variable across cells were removed using Caret v6.0-86 package. QC-ed cells were then segregated in two groups based on the number of neighbors: isolated cells having no neighbors and colony cells having one or more neighbors. Individual morphology traits were then summarized to well level measurement by averaging them across all cells per well, resulting in a well by trait matrix. Following this, each morphology trait was gaussianized across all 7 plates using inverse normal transformation (INT) method.

#### Selection of traits for association analysis

A set of morphology traits for association analysis (with both common variants and rare variant burden) was selected by considering their pairwise correlation across colony and isolate cells in the following steps: Step 1. Calculate Pearson correlation matrix for colony and isolate cells at donor level (total 2 correlation matrices). Step 2. Identify that single trait having the *Pearson r* >= 0.9 with the largest number of other traits across both correlation matrices. We specifically chose *Pearson r* >= 0.9 as cutoff here because most traits (93.7% and 91.2% traits in colony and isolated cells, respectively) had a correlation *Pearson* >= 0.9 with at-least one other trait (Fig S7). Step 3. Include that individual trait for association analysis. Remove it and other traits having *Pearson* >= 0.9 with it from correlation matrices. Step 4. Repeat step 1 to 3 until there are no more traits to include in association analysis.

#### Whole genome sequencing (WGS), variant calling and genes to test

DNA was obtained from cell line pellets with the Qiagen Quick-Start DNeasy Blood and Tissue Kit (cat. no. 69506). DNA samples were submitted to the Genomics Platform at the Broad Institute of MIT and Harvard. Whole genome sequencing (30x) was performed for all individuals (*n*=297) at the Broad Institute Genomics Platform using Illumina Nextera library preparation, quality control, and sequencing on the Illumina HiSeq 2500 platform. Raw sequences were QC-ed and sequencing reads (150 bp, paired-end) were aligned to the hg38 reference genome using the BWA alignment program. Variants were called and annotated (VQSLOD filter) using HapMap reference.

#### WGS data quality control for common variant association analysis

The QC-ed WGS VCF file was processed using plink v1.90b3 to remove sex chromosomes, multi-allelic variants, variants with duplicated positions, and small insertions and deletions larger than 5bp. Of 38,239,223 variants loaded from the VCF file, 33,348,914 passed these filters. Donor-level genotype missingness rates were checked to exclude donors with genotype missingness rates > 10%. All 297 individuals passed this filter. Finally, variants with minor allele frequency (MAF) < 5%, missingness > 5%, and Hardy-Weinberg equilibrium p-value < 10^-5^ were excluded, following which, 7,020,633 remained for common variant association analysis.

#### Principal components analysis (PCA)

Plink v1.90b3 was used on common (MAF > 5%) and post-QC variants to remove regions with known long-range linkage disequilibrium (LD) and variants in high LD (r2 > 0.1 in a window of 50 kb and a sliding window of 10 kb) (Price A. L. Am. J. Hum. Genetics 2008). The remaining 291,493 variants were loaded to GCTA v1.91.1 to generate a genetic relatedness matrix (GRM) using the --make-grm command with default options. The resulting GRM was used to generate 20 PCs using GCTA v1.91.1 --pca command with default options.

#### Variance component analysis

Variance component of fixed (cell neighbor density and donor’s age) and random effects (iPSC source tissue, cell line ID, plate and well of imaging, donor’s sex, and disease status) was estimated for selected traits using linear mixed model (lmer function in lmertest package). We included the first 4 PCs derived from genetic variation, corresponding to the elbow in scree plot, for ancestry/population stratification. The p-value of each factor was Bonferroni corrected for the number of traits.

#### Common variant association analysis

The linear regression framework implemented in GCTA v1.91.1 (--fastGWA-lr command) was used to test the association of common (MAF > 5%), post-QC variants with 246 post-QC, INT traits that were described above. Like the rare variant association analysis, plate and sex were included as categorical and four genotyping PCs, number of cell neighbors (for cells in colony) and the edge variable were included as quantitative variables in the model. Associations were considered statistically significant if they passed the genome-wide significance threshold for 246 tests (*P* < 5×10^-8^/246).

#### Rare variant burden test

The variants that were autosomal, passed the VQSLOD filter and called in >95% individuals were retained and annotated for their functional effect using SnpEff v5.0. To perform the rare variant burden test, the variants which were autosomal, passed the VQSLOD filter and called in >95% individuals and had MAF < 1% were retained. These variants were annotated for their functional effect using SnpEff v5.0. After annotation, those variants were kept which resided in the protein-coding region and had high or moderate effects on encoded protein. For each gene, multiple rare variants were grouped and coded as present or absent. The association between individual morphology traits and the presence of rare variants in a gene was investigated using linear regression models. The p-values of associations were corrected for both the number of tested traits and the number of genes using Bonferroni correction method.

#### CRISPRi sgRNA design, cloning, and virus production

To functionally validate the rare-variant burden associations, we designed sgRNAs targeting the transcriptional start site (TSS) for each gene using CRISPick software (Doench, 2016, Sanson, 2018). sgRNA oligonucleotides were cloned into the CROPseq vector using a Golden Gate cloning protocol (Juong, 2017). To validate sequence insertion, DNA plasmids were sequenced by a 3rd party provider. Plasmids with successful insertion were packaged for lentivirus generation using *Trans*IT-293 reagent (Mirus Bio cat#: MIR 2704) and packaging plasmids VSV-G and DVPR (need to confirm these). HEK239T (ATCC cat#: CRL-3216) cells were transfected with sgRNA packaging plasmid and incubated for 48hrs. HEK239T media supernatant was collected, and lentivirus was concentrated using LENTI-X concentrator (Takara) per manufacturer’s instructions. Virus supernatant was then aliquoted and stored at −80C.

#### sgRNA transduction in dCas9-iPSCs

An iPSC line, WTC11_TO-NGN2_dCas9-BFP-KRAB (gift from Michael Ward), was seeded at 250k cells per well in a 12 well plate and 50ul of sgRNA lentivirus was added to each designated well. The following day, 1mL of mTeSR1 complete media was added on top of the existing media. 48hrs post transduction, cells underwent a full media change with the addition of 1 ug/ml puromycin (Sigma Aldrich cat#: P8833) for chemical selection of cells which did not uptake the sgRNAs. Puromycin is supplemented in the feeding media for the duration the cell line is in culture to avoid uninfected cells from populating the dish.

#### qPCR analysis

RNA isolation was performed with the Direct-Zol RNA miniprep kit (ZYMO: cat# R2051) according to the manufacturer’s instructions. To prevent DNA contamination, RNA was treated with DNase I (ZYMO: cat# R2051). The yield of RNA was determined with a Denovix DS-11 Series Spectrophotometer (Denovix). 200 ng of RNA was reverse transcribed with the iScript cDNA Synthesis Kit (Bio-Rad, cat# 1708890). For all analyses, RT–qPCR was carried out with iQ SYBR Green Supermix (Bio-Rad, cat# 1708880) and specific primers for each gene (listed below) with a CFX384 Touch Real-Time PCR Detection System (Bio-Rad). Target genes were normalized to the geometric mean of control genes, *RPL10* and *GAPDH*, and relative expression compared to the mean Ct values for non-targeting control sgRNAs and gene targeting sgRNAs, respectively.

The following primers were used:

WASF2_forward 5’-TAGTAACGAGGAACATCGAGCC-3’

WASF2_reverse 5’-AAGGGAGCTTACCCGAGAGG-3’

PRLR_forward 5’-TCTCCACCTACCCTGATTGAC-3’

PRLR_reverse 5’-CGAACCTGGACAAGGTATTTCTG-3’

TSPAN15_forward 5’-TCCCTCCGTGACAACCTGTA-3’

TSPAN15_reverse 5’-CCGCCACAGCACTTGAACT-3’

RPL10_forward 5’-GCCGTACCCAAAGTCTCGC-3’

RPL10_reverse 5’-CACAAAGCGGAAACTCATCCA-3’

GAPDH_forward 5’-GGAGCGAGATCCCTCCAAAAT-3’

GAPDH_reverse 5’-GGCTGTTGTCATACTTCTCATGG-3’

## Data Availability

Images and preprocessed profiles that are augmented with gene and compound annotation are available in the Cell Painting Gallery on the Registry of Open Data on AWS (https://registry.opendata.aws/cellpainting-gallery/) as dataset ‘cpg0022-cmqtl’ at no cost and no need for registration. Whole genome sequencing for cell lines used in this study are hosted on Terra https://app.terra.bio/#workspaces/anvil-datastorage/AnVIL_NIMH_Broad_ConvergentNeuro_McCarroll_Eggan_CIRM_GRU_WGS

## Code Availability

Source code to reproduce and build upon the presented results is available at https://github.com/broadinstitute/cmQTL

## Acknowledgements

We thank members of the Nehme, Carpenter-Singh, and Raychaudhuri labs for insightful discussions and critical reading of the manuscript, and Anna Neumann for project management. This work was supported by a Broad Institute Variant to Function (V2F) Initiative grant, the National Institutes of Health (NIMH U01 MH115727 to RN/KE/SAM and NIGMS MIRA R35 GM122547 to AEC), the Chan Zuckerberg Initiative DAF, an advised fund of the Silicon Valley Community Foundation (grant number 2020-225720 to BAC), as well as the Stanley Center for Psychiatric Research at the Broad Institute. The authors also gratefully acknowledge the use of the PerkinElmer Opera Phenix High-Content/High-Throughput imaging system at the Broad Institute, funded by the S10 Grant NIH OD-026839-01.

## Author contributions

M.T., J.A., S.A., A.E.C., S.S., R.N., and S.R., conceived the work and wrote the manuscript. R. N. obtained cell lines from CIRM and coordinated the sequencing. M.T., with support from E.P. and D.L., performed cell culture data generation and CRISPRi knockdown validation. J.A., S. A., performed genetic association analyses. B.A.C., S.S., G.W., M.H., E.W., processed and analyzed Cell Painting data. A.E.C., S.S., R.N., and S.R. supervised the work and analyses.

## Competing interests

The authors declare no competing interests.

## Supplemental Figures

**Fig S1.**
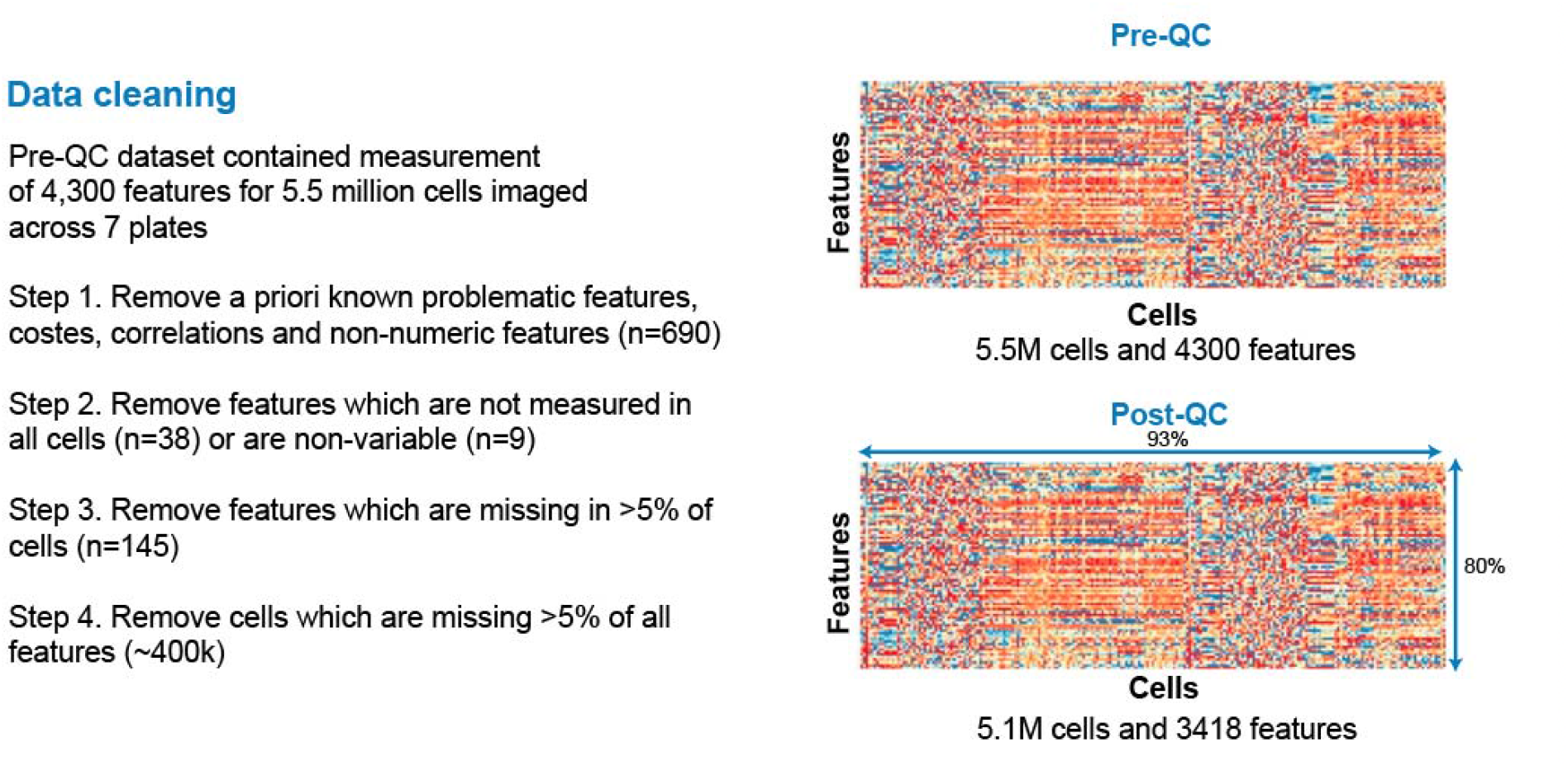
Data filtering and quality control for the traits measured across iPSC cells. A total of 4318 cell morphology traits were quantified across all 5.5 million iPSCs cells from 297 donors. Morphology traits a priori known to be problematic, not measured across all cells or non-variable across cells were removed. Also, cells missing measurement for >5% of traits were removed, yielding 3418 traits across 5.1 million cells.

**Fig S2.**
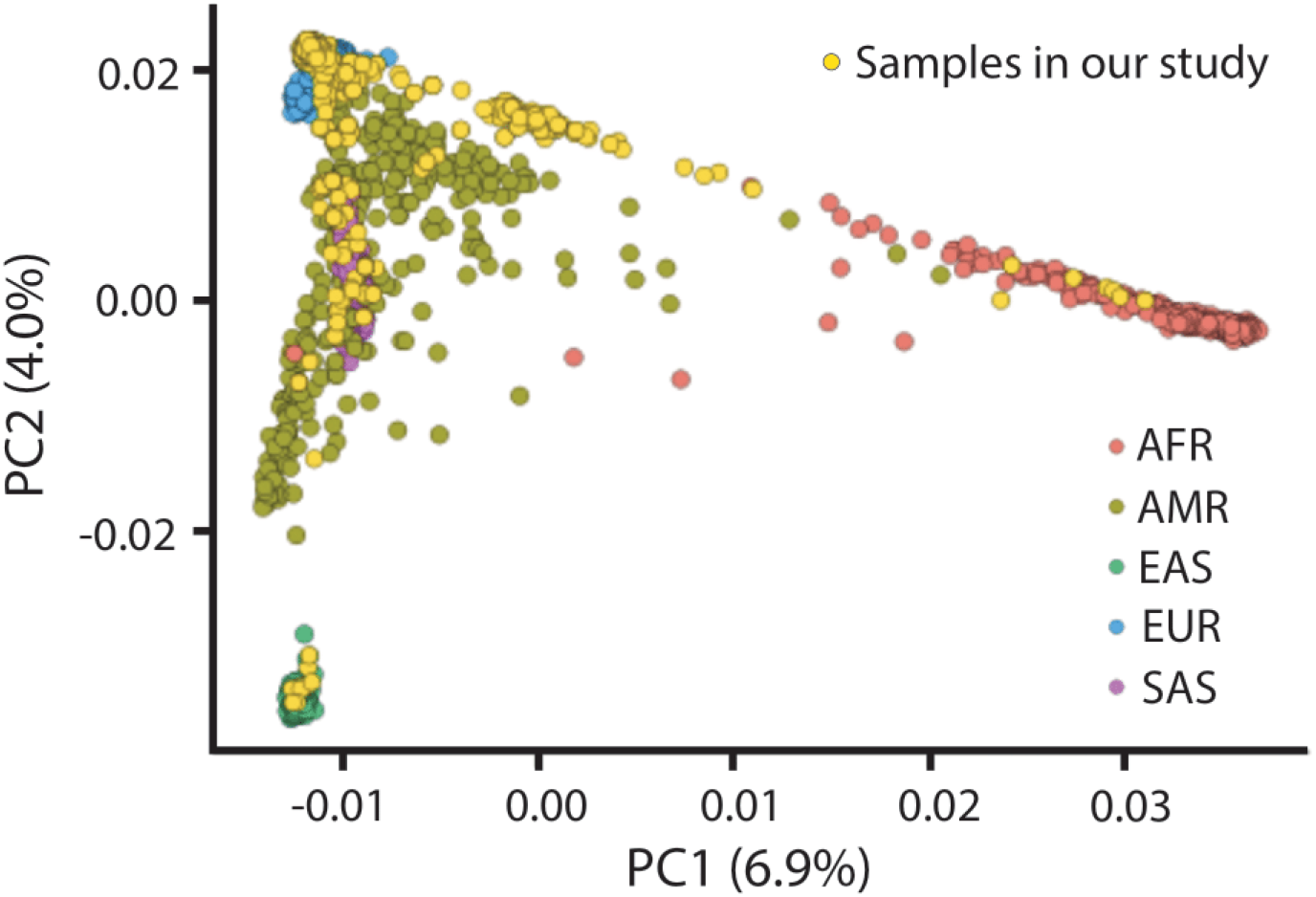
Principal component analysis (PCA) of donors. Distribution of 297 donors (yellow dots) laid over individuals from 1K genomes on PC1 and PC2 calculated from common variants (maf > 5%). Of 297 donors, 207 self-reported their ancestry as European.

**Fig S3.**
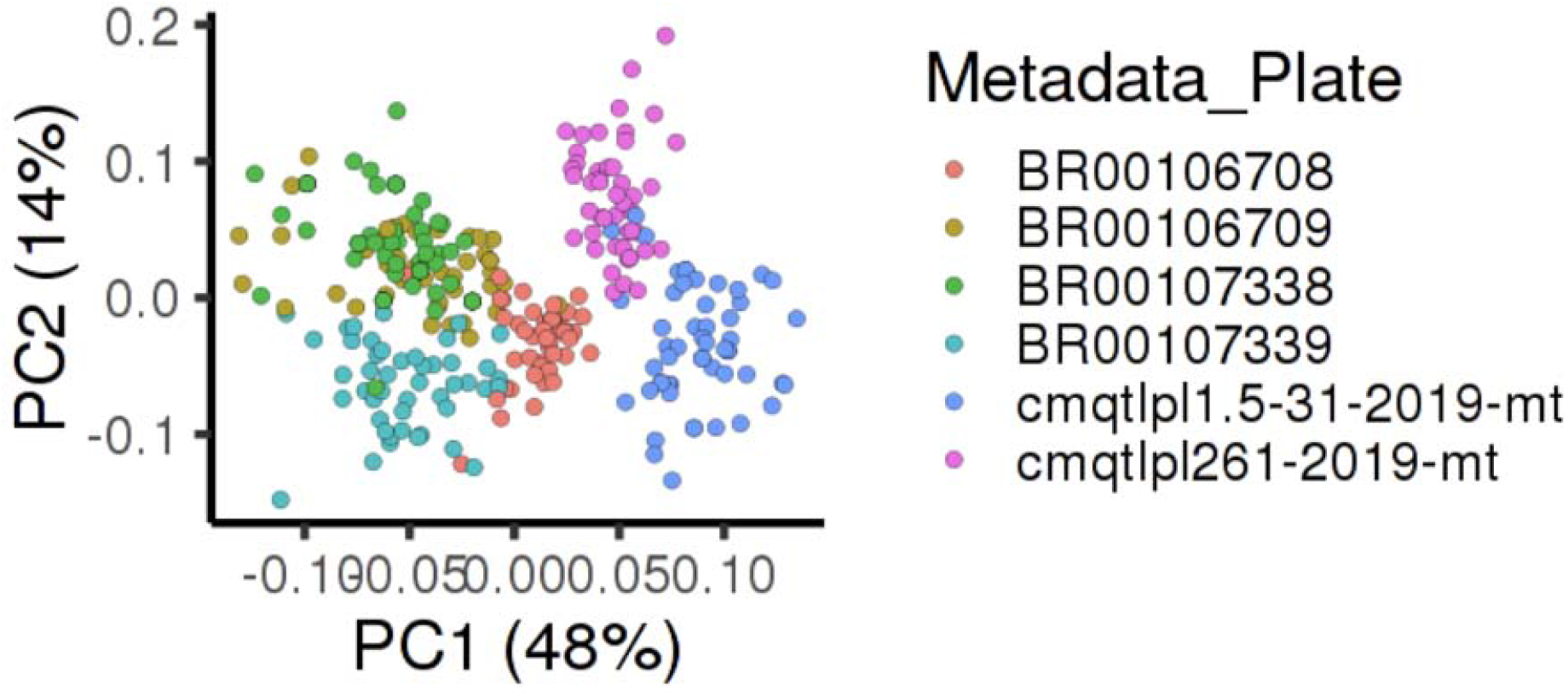
Batch effects in measurement of morphology traits. Distribution of 297 donors on PC1 and PC2 calculated from morphology traits (n=3418) color by 7 plates on which iPSCs from donors were imaged, showing the batch (plate) effect in the measurement of morphology traits.

**Fig S4.**
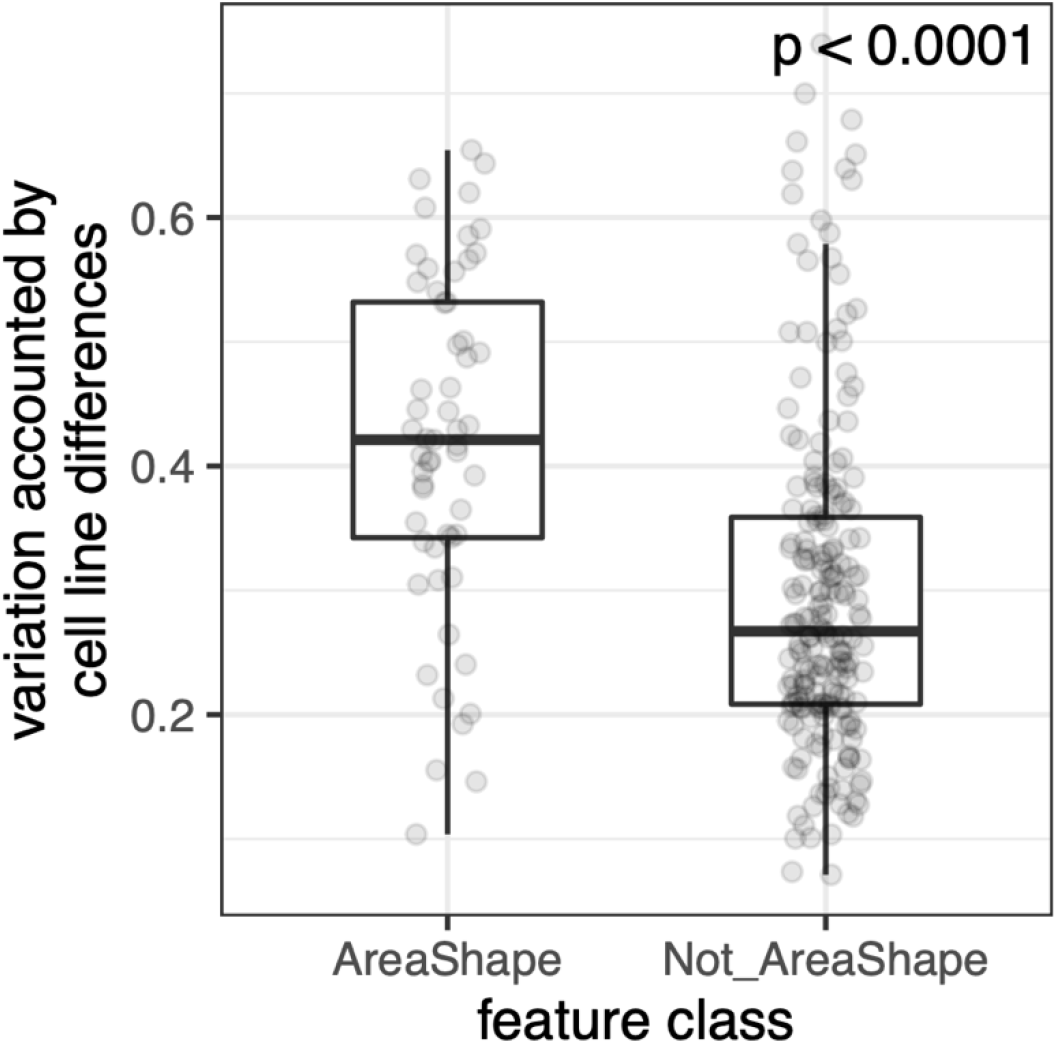
Variation in traits explained by genetic difference among donors. The comparison of variation explained by genetic difference among donors in traits belonging to Area and Shape category and other categories. P-value from Wilcoxon rank sum test is shown.

**Fig S5.**
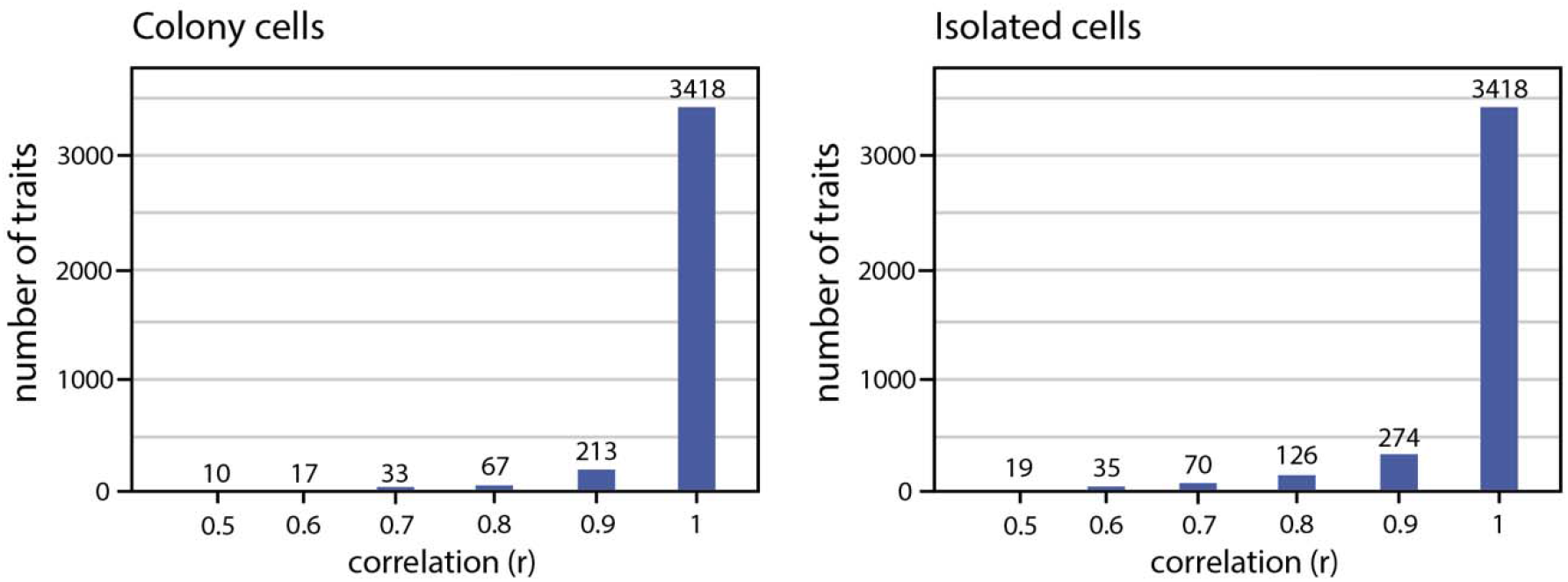
Correlation among morphology traits. The number of traits having correlation (Pearson r) of up to 0.5, 0.6, 0.7, 0.8, 0.9 and 1 (on x-axis) with at-least one other trait is shown for cells in colonies and cells which are isolated.

**Fig S6.**
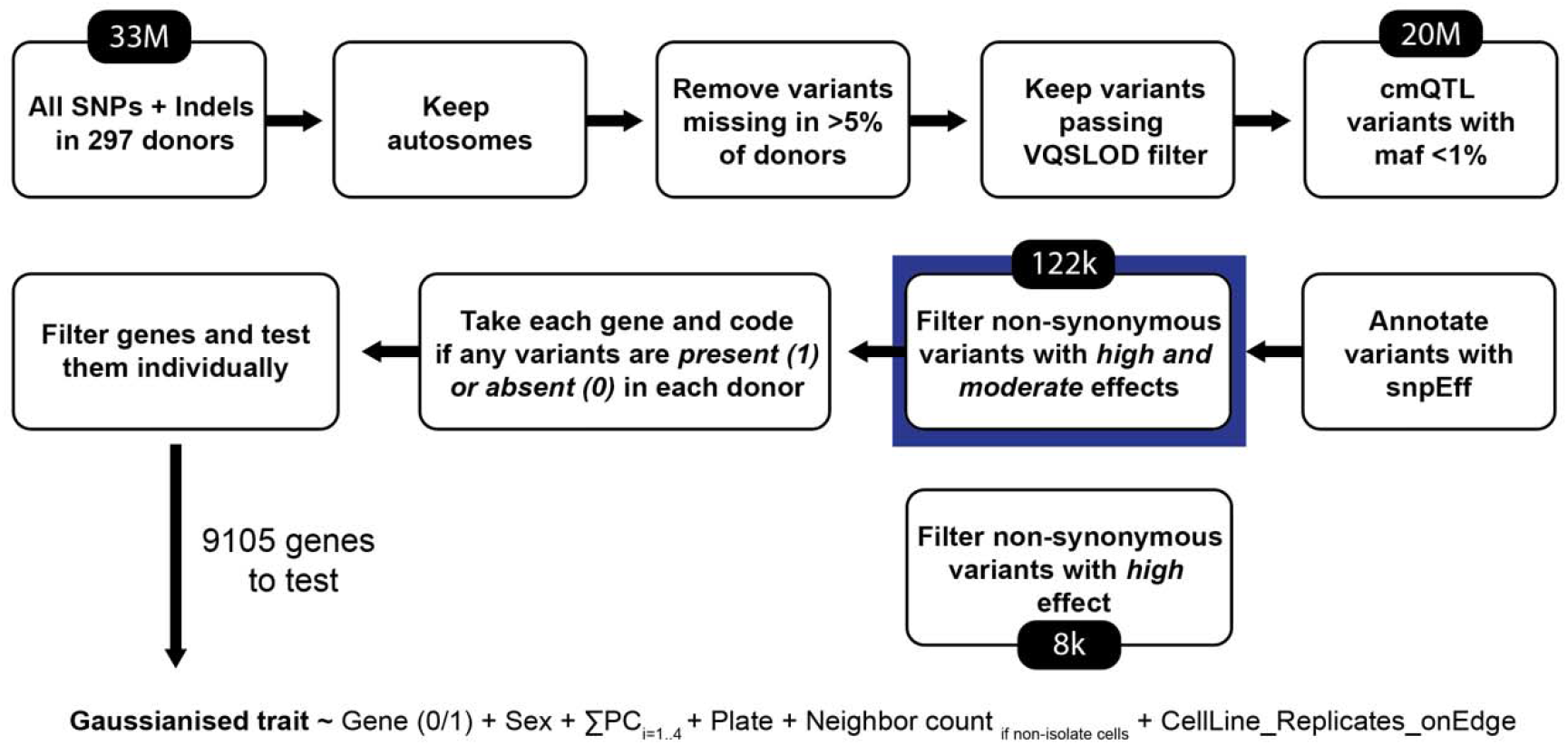
Rare variant workflow. Step by step workflow for QC and selection of rare variants for the association analyses.

**Fig S7.**
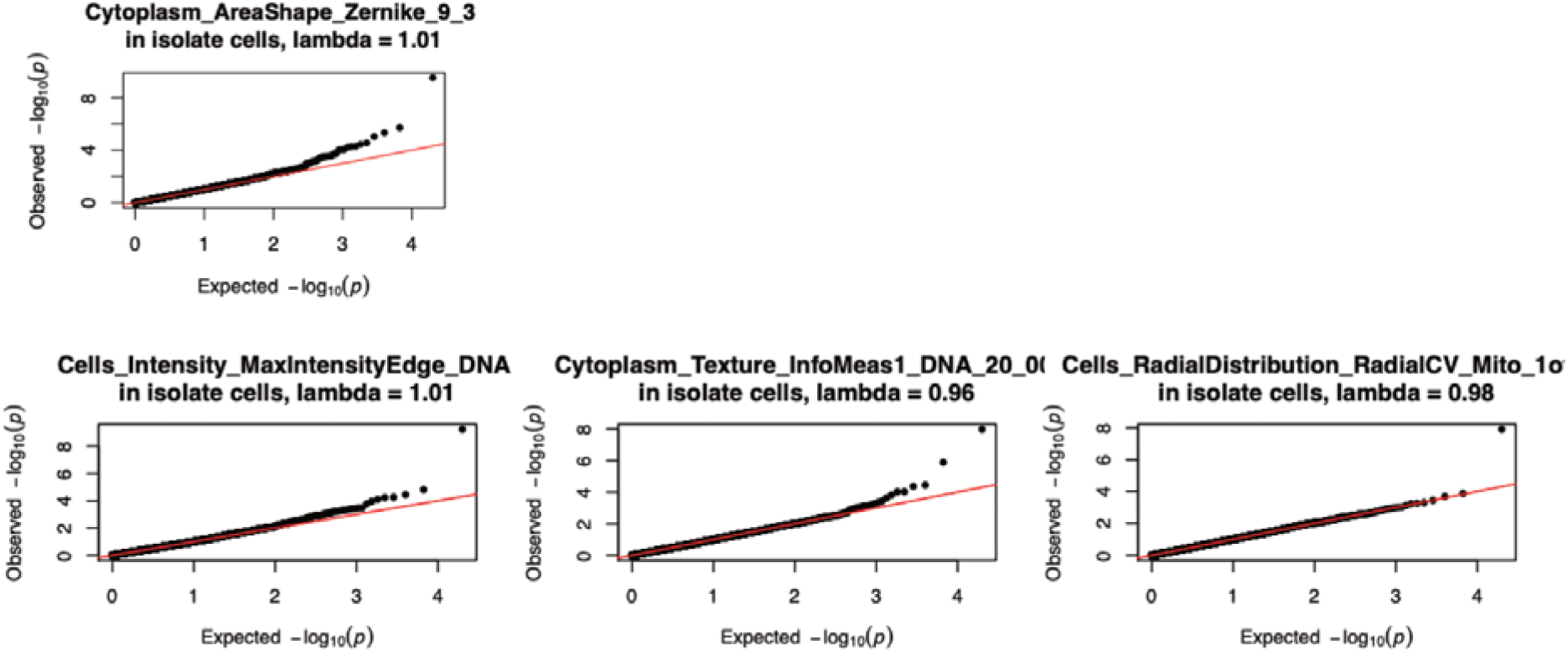
Quantile-quantile (QQ) plots for associated traits. QQ plots show the distribution of expected and observed p-value of association with all tested genes for 4 morphology traits. Each dot is a tested gene. Lambda statistic (λ), a measure of inflation in observed p-values, is shown.

**Fig S8.**
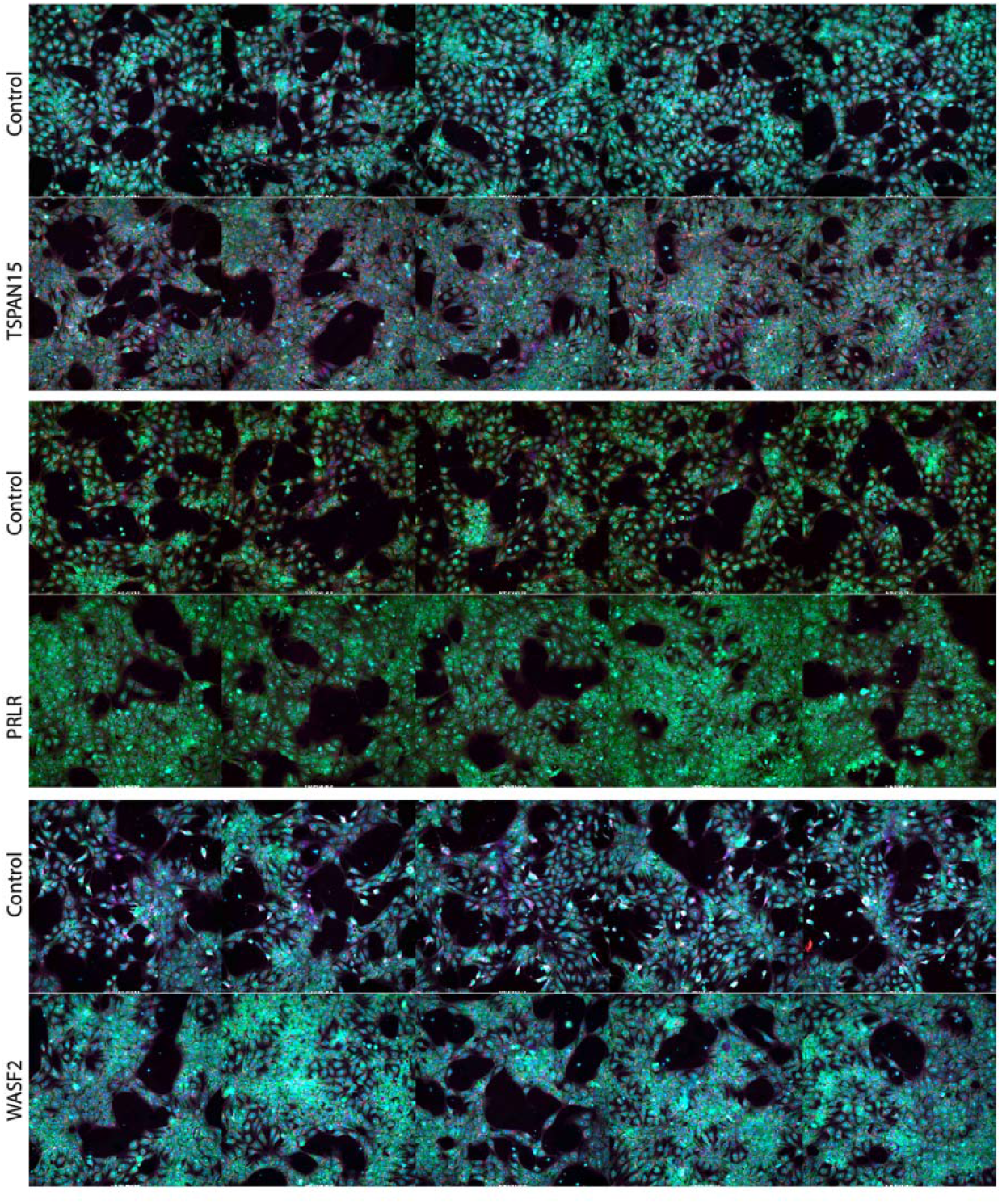
Representative images from cell lines with rare variants in WASF2, PRLR, and TSPAN15. Randomly selected representative images from wells containing cell lines harboring rare variants in *WASF2, PRLR*, and *TSPAN15* compared to reference cell lines with no detected variants.

**Fig S9.**
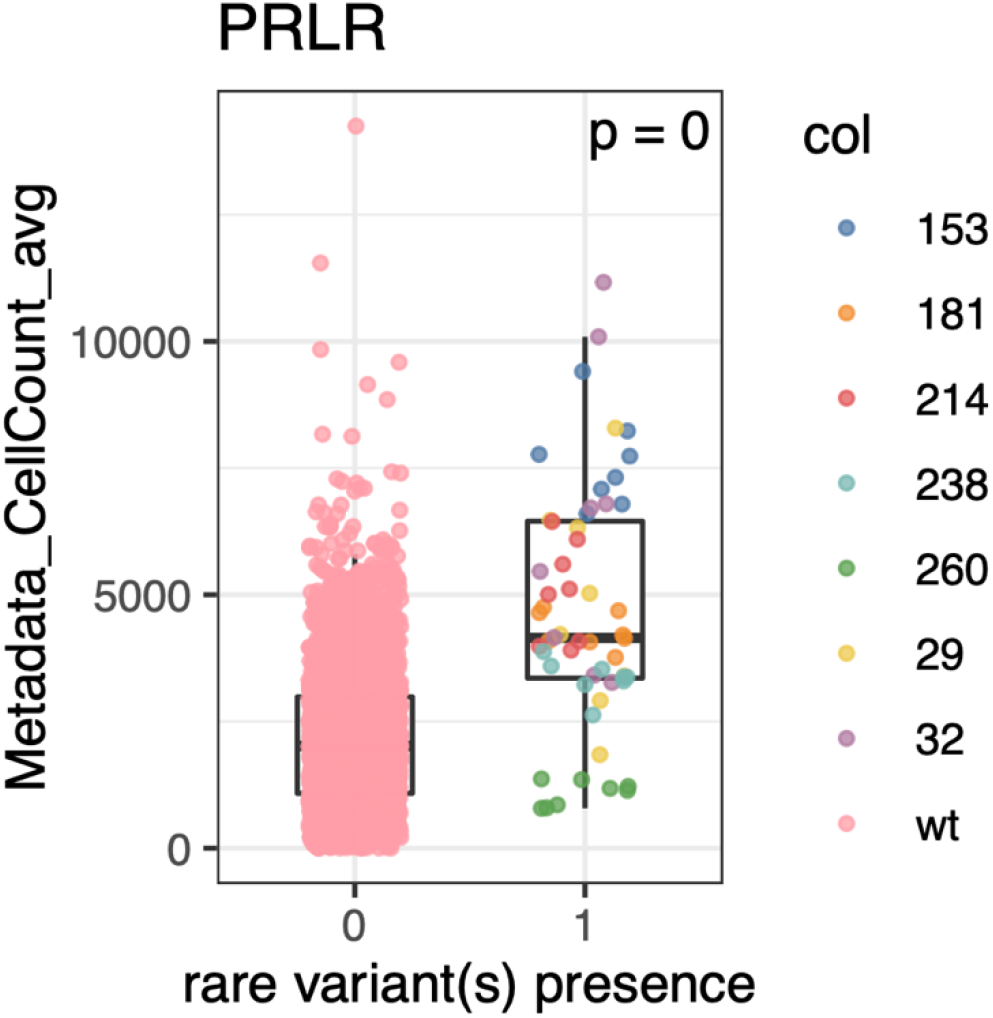
Cell count for *PRLR* cell lines compared to others. Boxplots displaying per well cell count between cell lines harboring rare variants in *PRLR* compared to reference cell lines

**Fig S10.**
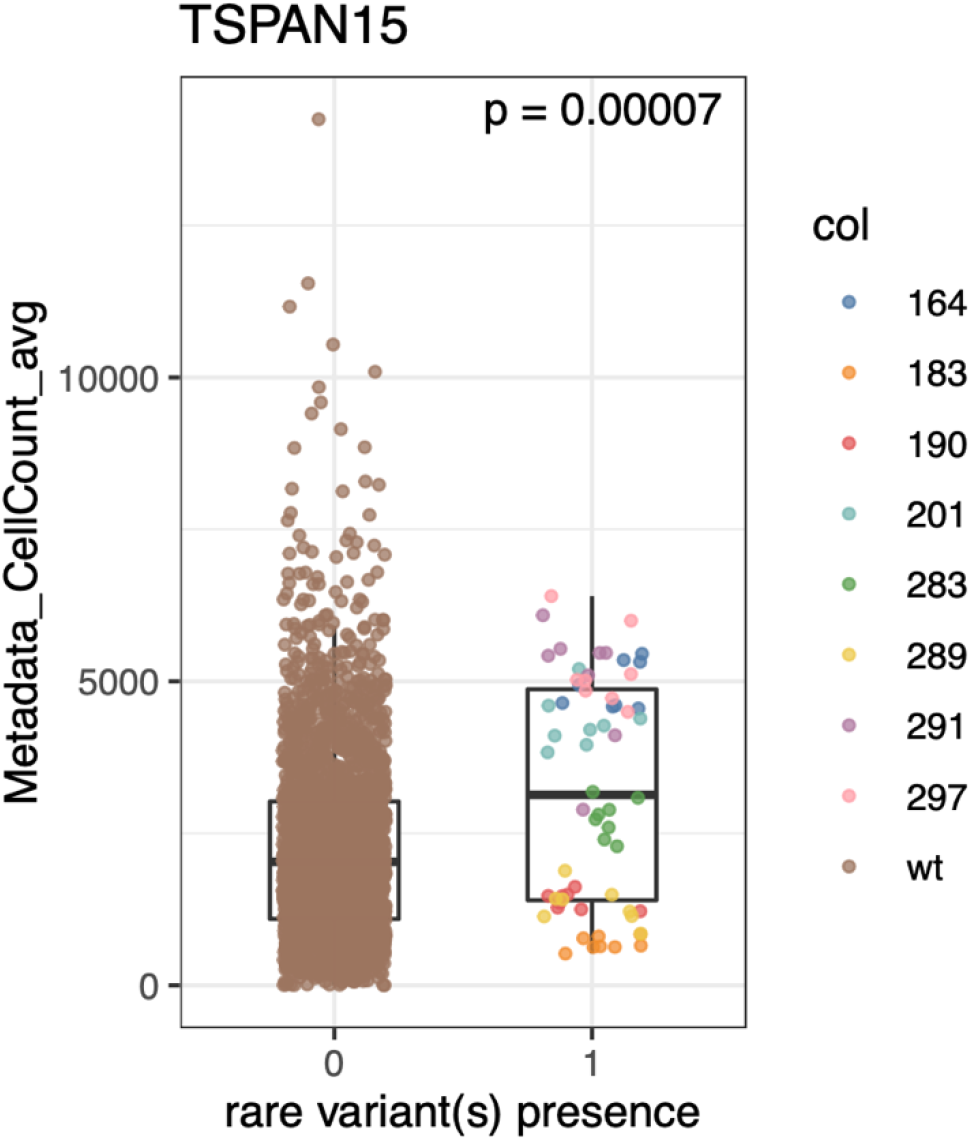
Cell count for *TSPAN15* cell lines compared to others. Boxplots displaying per well cell count between cell lines harboring rare variants in *TSPAN15* compared to reference cell lines

**Fig S11.**
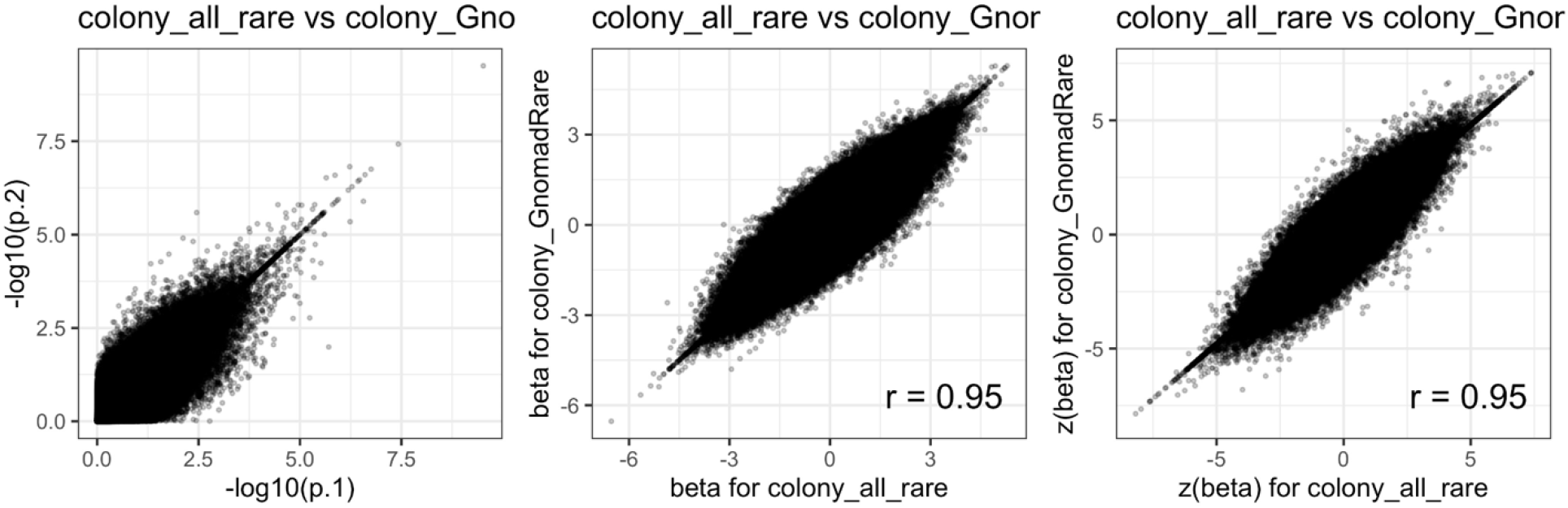
Associations using rare variants present in Gnomad. Comparison of p-value and z-score of effect size (beta) of associations between individual morphology traits and rare variant burden in individual genes using all rare variants in our dataset and those rare variants (out of all) which are also present in Gnomad dataset is shown for colony cells (A) and isolated cells (B). The orange colored dots represent significant associations from Fig 3A where we used all rare variants. Pearson r is shown.

**Fig S12.**
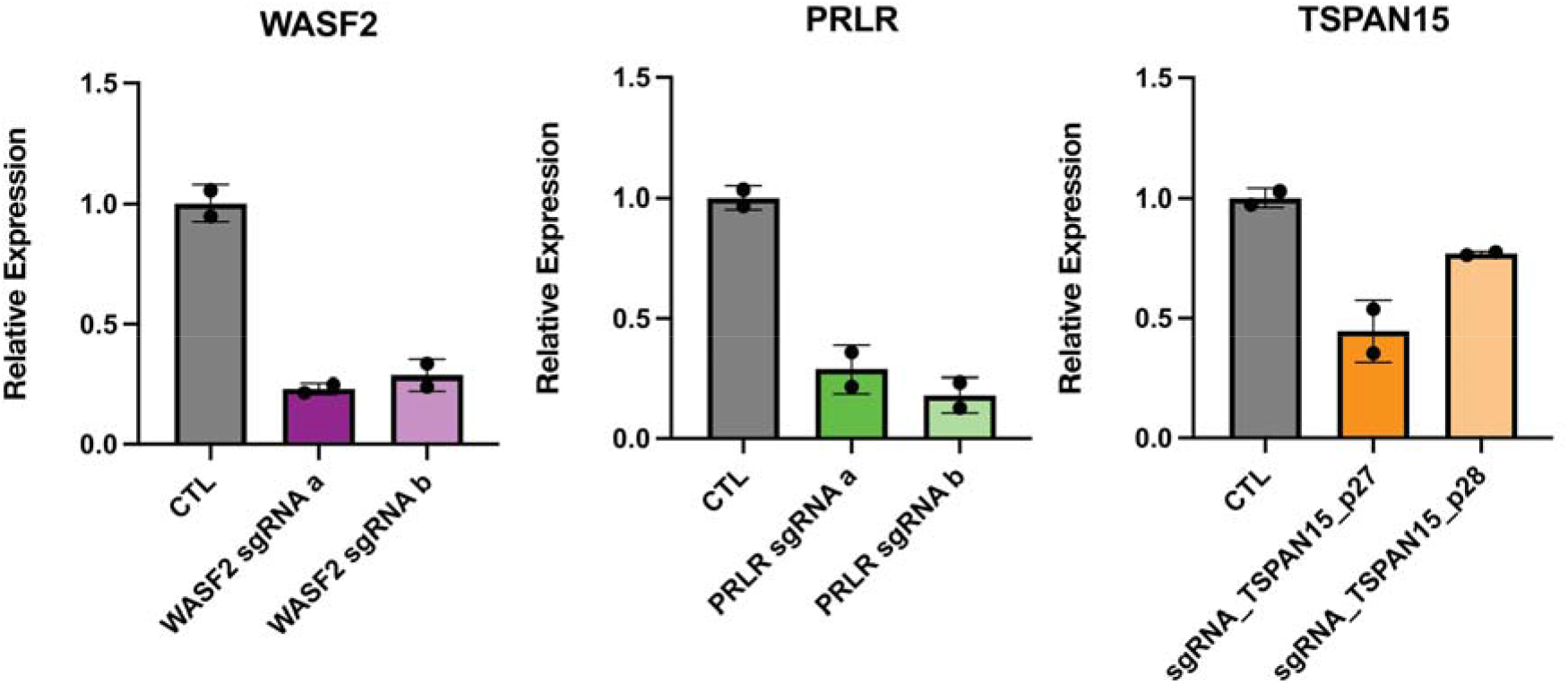
qPCR knockdown of rare-variant associations using CRISPR interference. Relative expression of sgRNA target genes compared to *GAPDH* and *RPL10* between iPSCs transfected with gene targeting sgRNAs and non-targeting control sgRNAs.

## Supplemental Tables

Table S1. Cell line metadata

Demographic characteristics for all 297 iPSC lines used in this study.

Table S2. All morphological traits

All 3418 morphological traits which passed QC

Table S3. Composite morphological traits

246 traits which were used for the association tests.

Table S4. Morphology trait associations with rare variant burden in *WASF2*

Morphological traits which meet nominal significance with association to rare variant burden in *WASF2*.

Table S5. Morphology trait associations with rare variant burden in *PRLR*

Morphological traits which meet nominal significance with association to rare variant burden in *PRLR*.

Table S6. Morphological traits with suggestive evidence of association with rare variants in our study

Morphological traits which show suggestive evidence for association with rare variants in our study.

Table S7. CRISPRi sgRNA sequences

Oligonucleotide sequences for all sgRNAs used in this study.

Table S8. Morphological traits with suggestive evidence for association to common variants in our study.

Morphological traits which show suggestive evidence for association with common variants in our study.

## Notes

https://registry.opendata.aws/cellpainting-gallery/

### Competing Interest Statement

The authors have declared no competing interest.

